# DeepLabCut-based Behavioural and Posture Analysis in a Cricket

**DOI:** 10.1101/2023.11.14.567118

**Authors:** Shota Hayakawa, Kosuke Kataoka, Masanobu Yamamoto, Toru Asahi, Takeshi Suzuki

## Abstract

Circadian rhythms are indispensable intrinsic programs that regulate the daily rhythmicity of physiological processes, such as feeding and sleep. The cricket has been employed as a model organism for understanding the neural mechanisms underlying circadian rhythms in insects. However, previous studies measuring rhythm-controlled behaviours only analysed locomotive activity using seesaw-type and infrared sensor-based actometers. Meanwhile, advances in deep learning techniques have made it possible to analyse animal behaviour and posture using software that is devoid of human bias and does not require physical tagging of individual animals. Here, we present a system that can simultaneously quantify multiple behaviours in individual crickets, such as locomotor activity, feeding, and sleep-like states, in the long-term, using DeepLabCut, a supervised machine learning-based software for body keypoints labeling. Our system successfully labelled the six body parts of a single cricket with a high level of confidence and produced reliable data showing the diurnal rhythms of multiple behaviours. Our system also enabled the estimation of sleep-like states by focusing on posture, instead of immobility time, which is a conventional parameter. We anticipate that this system will provide an opportunity for simultaneous and automatic prediction of cricket behaviour and posture, facilitating the study of circadian rhythms.

## Introduction

Circadian rhythms are endogenous oscillations observed in most animals. Circadian rhythms have a free-running period of approximately 24 h, entrained by daily environmental cues, such as light and temperature. In insects, circadian rhythms control the daily rhythmicity of physiological and developmental processes, such as hatching, pupation, and moulting, as well as behaviour (*e.g.* locomotion, feeding, sleep, mating, and oviposition) (Saunders, 2002).

Recent advances in computer vision and deep learning have led to several open-source programs and commercial systems that accurately, rapidly, and robustly measure animal behaviour. Videography is the most common and widely used method for measuring animal behaviour because it is non-invasive and provides high-resolution behavioural observations. Although the extraction of animal behaviour from videos poses a computational challenge, recent developments in computer vision have offered numerous algorithms for processing image data. In particular, supervised machine learning methods that automatically label and track “keypoints” on an animal’s body can identify complex behavioural changes over time. These resources not only overcome the enormous time and effort involved in manually identifying behavioural traits but also eliminate human biases and oversights during annotation. This allows us to capture multiple behaviours and behavioural action sequences in a consistent and objective manner (Anderson and Perona, 2014), offering the potential to track a variety of behaviours under circadian control.

The above-described technological advancements facilitate further research into sleep, defined by the behavioural characteristics of invertebrates. The sleep-like state (SLS) is generally defined by the following five features: (i) increased arousal threshold, (ii) reversibility, (iii) homeostasis, (iv) decreased movement, and (v) a specific sleep posture (Campbell and Tobler, 1984). The latter is accompanied by reduced muscular activity, which has been observed in nearly all animals, from mammals to arthropods (sloths: De Moura Filho et al., 1983; horses: Dallaire, 1986; bees: Kaiser, 1988; cockroaches: Tobler and Neuner-Jehle, 1992; nematodes: Schwarz et al., 2012). For fruit flies, a behavioural measurement system, called the Drosophila Activity Monitor System (DAMS), was previously utilised to define SLS as continuous immobility lasting for more than 5 min (Shaw et al., 2000; Hendricks et al., 2000). Continuous immobility time has been used as an evaluation metric for the SLS of insects (Ajayi et al., 2022).

Crickets have served as a useful model animal for understanding the neural mechanisms underlying circadian rhythms in insects because they have relatively large nervous systems with identifiable neurons (Cymborowski, 1973; Abe et al., 1997; Tomioka and Abdelsalam, 2004; Tomioka, 2014). Crickets are suitable for manipulation through reverse genetic methods, such as RNA interference and genome editing (Mito et al., 2011; Watanabe et al., 2012). In the context of circadian rhythms, crickets have been suggested to exhibit a unique clock oscillation mechanism in which both the *cycle* and *clock* genes are rhythmically regulated (Uryu et al., 2013). Further, considerable advances have been made in cricket genomics (*e.g.*, *Laupala kohalensis*: Blankers et al., 2018; *Teleogryllus oceanicus*: Pascoal et al., 2019; *T. occipitalis*: Kataoka et al., 2020; *Gryllus bimaculatus*: Ylla et al., 2021; *Acheta domesticus*: Dossey et al., 2023; *Apteronemobius asahinai*: Satoh et al., 2021). These have enabled a more comprehensive study of circadian rhythms from a molecular biological perspective (Kataoka et al., 2022). An in-depth analysis of the behavioural rhythms of crickets is expected to facilitate the identification of novel roles for genes that determine circadian rhythm-controlled physiological behaviour. Previous studies measuring the daily rhythms of behaviour in crickets only adopted conventional analytical methods, such as the seesaw-type actometer (Moriyama et al., 2008) and an infrared sensor-based actometer (Yamano et al., 2001). However, these systems cannot measure activities other than locomotor activity.

In this study, we propose a method for quantifying and analysing multiple behaviours controlled by circadian rhythms, including locomotor activity, feeding, and SLS, in a cricket. To this end, we applied the open-source software DeepLabCut (Mathis et al., 2018; Nath et al., 2019; Lauer et al., 2022), which is based on the Convolutional Neural Network (CNN) of supervised machine learning, to automatically label the keypoints of cricket bodies and derive quantitative data from the coordinates of body parts. Unlike fruit flies, crickets can be accurately labelled at key points on their body parts owing to their large body sizes. This means that the system can automatically acquire not only quantitative measurements of behavioural activities such as locomotion, feeding, and resting, but also more complex information such as posture. In light of this advantage, we attempted to measure SLS based on posture, in addition to immobility time. Multiple behaviours were automatically detected from long-term recordings using an automated analysis package that can be easily implemented for a variety of animals. Furthermore, we propose a new method that contributes to a multifaceted definition of SLS that does not rely on restricted criteria, such as immobility time.

## Results

### High-precision keypoint detection

We set up an imaging system controlled by Raspberry Pi for the long-term quantification of a variety of behaviours in six adult crickets (Fig 1). We imaged the crickets, each compartmented in an acrylic case, once per min under the 12h light 12h dark conditions (12:12 LD) for 14 days. The resulting serial images were cropped into individual compartments and converted into videos for DeepLabCut (Movie). In DeepLabCut, we manually labelled the keypoints of body parts (*i.e.* head, thorax, abdomen, abdominal tip, left hind leg, and right hind leg) in randomly extracted frames. A ResNet50 neural network was used to train the manually labelled images (training process). The resulting model was applied to test the newly extracted frames, and the labelling predicted by this model was manually corrected (refinement process). The training and refinement processes were repeated five times. The corrected labelling generated by the refinement process was used for subsequent training. Although the losses in our models did not converge close to zero in the first three training sessions, they converged near zero after the fourth session (Fig S1A). We used the average of the absolute Euclidean distances to compare the keypoints generated by the model with those labelled by humans. This distance was obtained by calculating the Euclidean distance between keypoints generated by the model and those labelled by humans, for every frame. We computed the training error generated during training and the test error generated during prediction using the model (Fig S1B). The training and test errors for the first model were 3.32 px and 5.59 px, respectively (Fig S1B). In the final model, the training and test errors were reduced to 1.61 px and 1.58 px, respectively, which are nearly equivalent (Fig S1B and S1C). These results suggest that accurate body part predictions were achieved in crickets after five rounds of training and correction of model prediction.

**Fig. 1.**
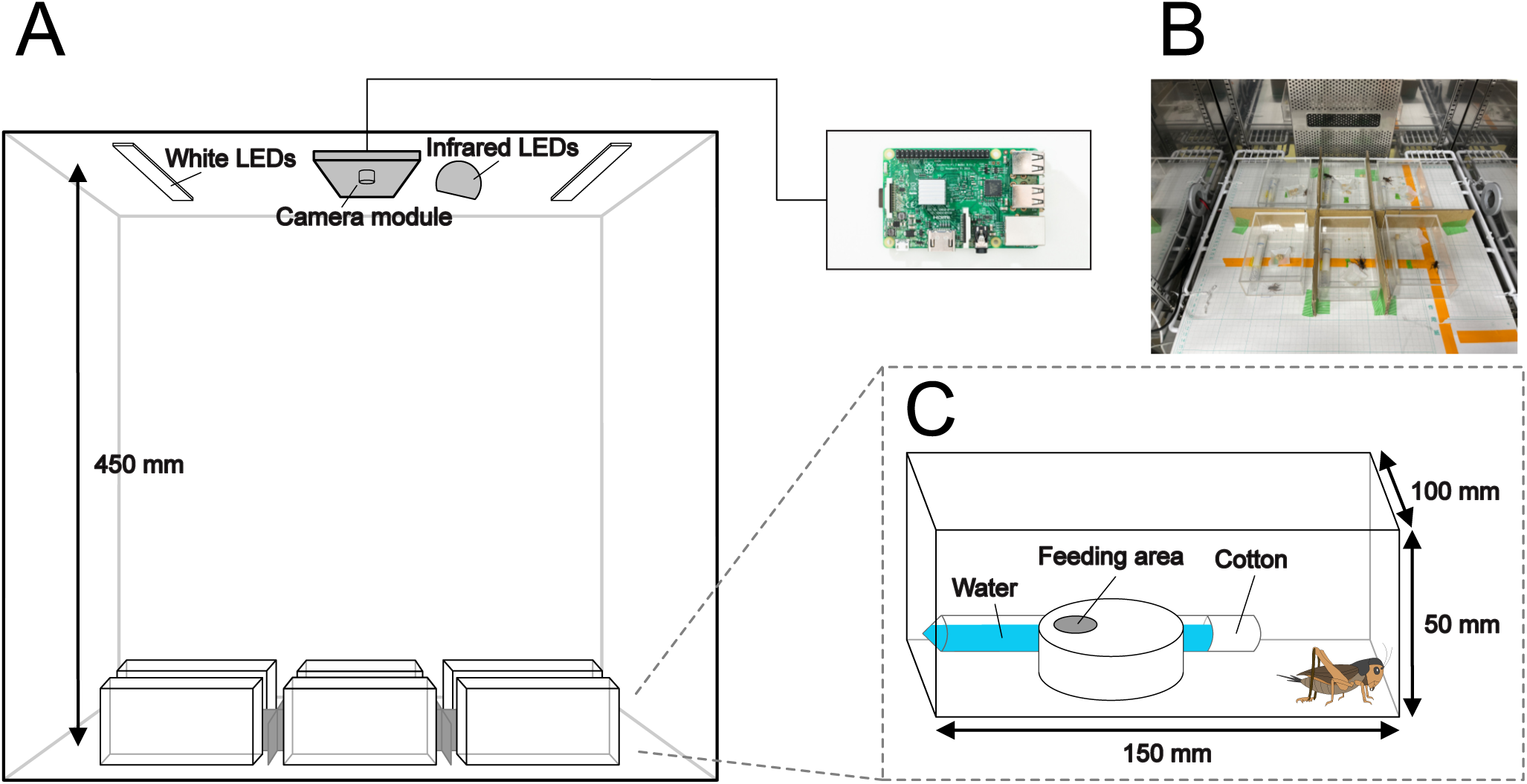
A schematic of the cricket behaviour-recording system. (A) An overview of the quantitative evaluation system for recording, through which we simultaneously recorded up to six crickets in one experiment by isolating individuals using acrylic cases (150 × 100 × 50 mm). The 12-h light/dark cycle (LD 12:12) was controlled by an electric time switch. The movement of crickets was recorded by a video camera, set at a height of 450 mm above the floor and connected to a Raspberry Pi 3. (B) A photograph depicting the specific recording environment. Sheets of paper with a height of 15 mm were placed between each acrylic, to make the individuals invisible to each other. (C) A schematic diagram of the acrylic case. The crickets were allowed free access to water and feed. The water source is the tube-stuffed cotton, and the feeding area consists of a section of the plastic case with holes.

Using the final model, we predicted six keypoints for crickets in all serial images over a 14-day period. To evaluate the labelling accuracy, the ratio of each keypoint with a likelihood greater than 0.95 was calculated for all frames (Fig S1D). In all individuals, accurate labelling was achieved in the head, thorax, abdomen, and abdominal tip in more than 90% of frames (96.39 ± 1.45% for head, 97.31 ± 1.42% for thorax, 97.27 ± 1.68% for abdomen, and 97.57 ± 0.83% for abdominal tip). In the left hind leg and right hind leg, there tended to be fewer frames with a likelihood greater than 0.95 than in the other four points (91.59 ± 1.70% for left hind leg, and 93.20 ± 1.34% for right hind leg). We also checked for differences in labelling accuracy between the dark and light periods. There was a trend toward reduced labelling accuracy during the dark period for all six key points (Fig S1E). The decrease in accuracy in the left and right hind legs was particularly remarkable (Fig S1E). These results were likely due to the reduced resolution of images acquired under infrared light conditions.

### Locomotor activity

Locomotor activity was recorded for the six crickets during exposure to 12:12 LD for 14 days. The criterion for judging locomotion was defined as whether the coordinates of the abdomen moved more than the body length (from the head to the abdominal tip) within a single frame (Fig 2). A double-plotted actogram was generated based on the data from all frames (Fig 3A). They showed a clear nocturnal rhythm, with weak activity that started several hours before turning off the lights also noted (Fig 3B, C), which is consistent with previous studies in other cricket species (*Acheta domestica*: Cymborowski, 1973; *Gryllodes sigillatus*: Abe et al., 1997; *Gryllus bimaculatus*: Tomioka and Abdelsalam, 2004). For most individuals, acrophases (circadian peak time) of the daily rhythms of locomotion were observed immediately after lights-off (Fig 3C, Fig S2A). The circadian rhythms of most animal locomotion were entrained to nearly 24 h (1440 min) by the 12:12 LD photocycle, as demonstrated by Chi-square and Lomb-Scargle periodograms (Fig S2B). These findings indicate that the quantification of locomotor activity using DeepLabCut produces reliable results comparable to those of other available methods.

**Fig. 2.**
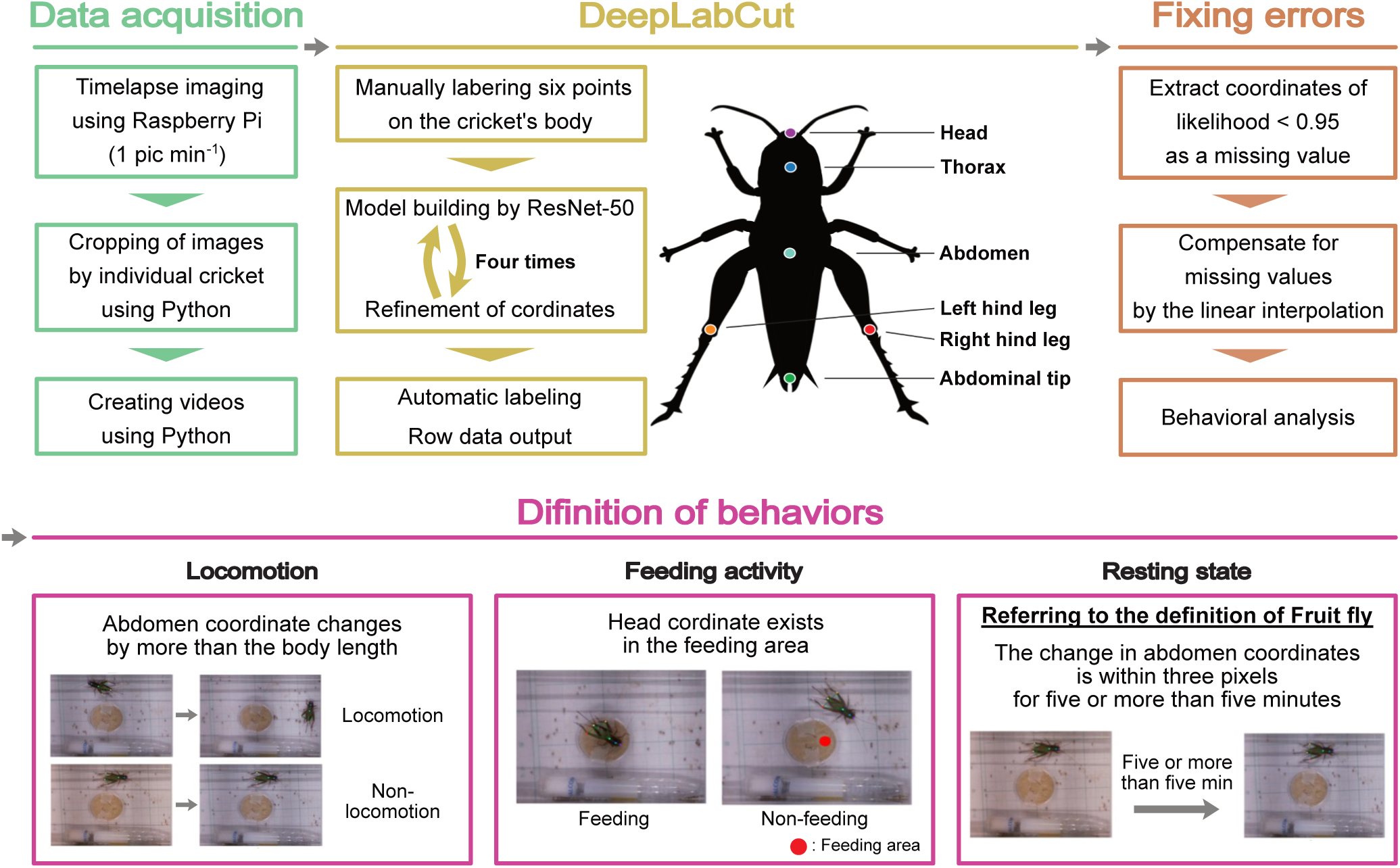
Outline view of the system. Scheme from image acquisition to behavioural quantification. First, image data were recorded with a camera module connected Raspberry Pi at an interval of one image per minute. Images were converted into videos, and a machine learning algorithm, ResNet-50 and DeepLabCut, was employed for training and refinement to construct a model, with six automatically labelled keypoints on crickets; head, thorax, abdomen, abdominal tip, left hind leg, and right hind leg. Subsequently, the coordinates of these labelled keypoints were extracted. After fixing errors, behaviours such as locomotion, feeding activity, and sleep-like state were defined based on the coordinates and behaviours quantitatively analysed across all frames over a period of 14 days.

**Fig. 3.**
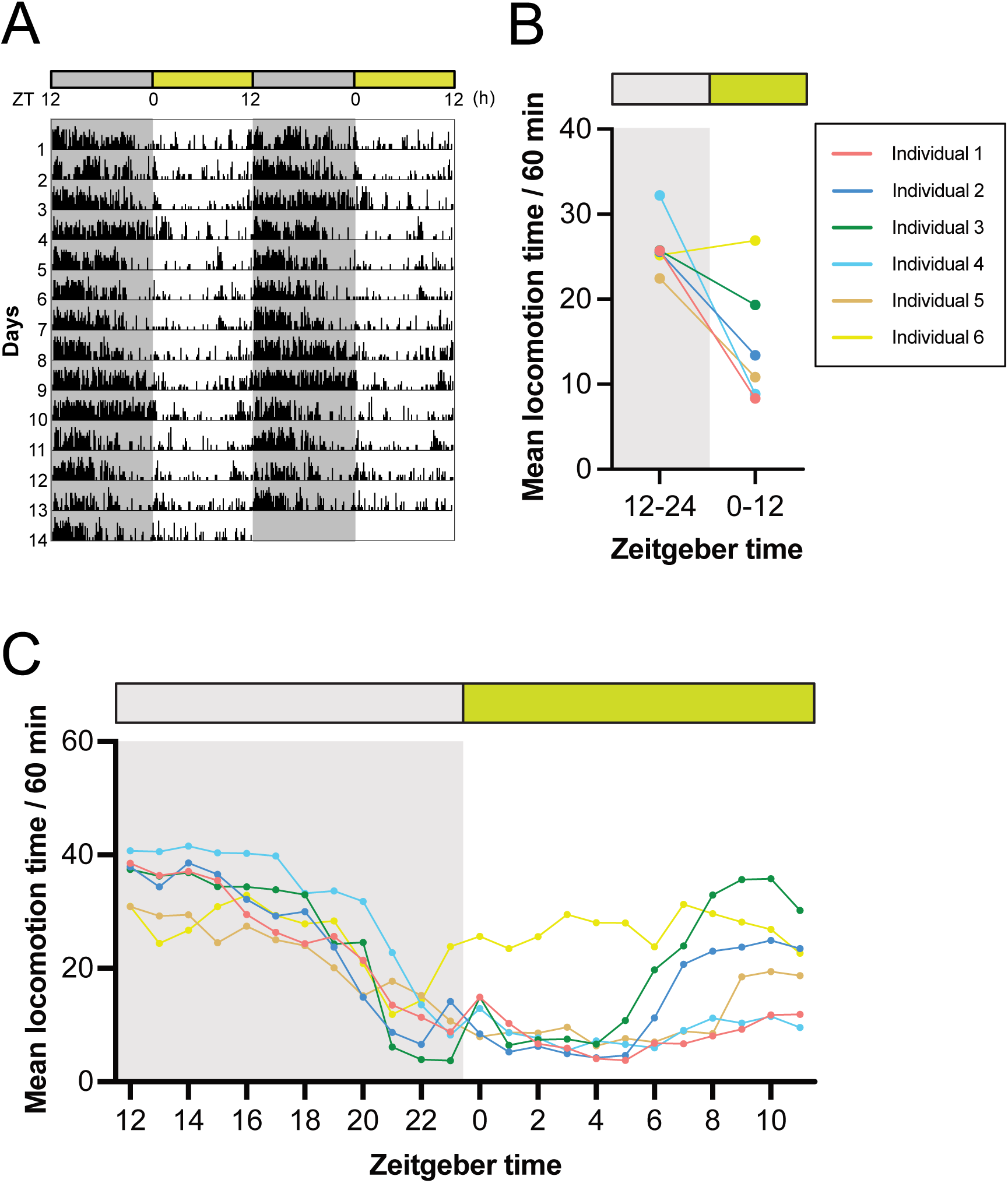
A quantitative analysis of locomotion of crickets. (A) The double-plotted actogram recorded under a 12-h light:dark (LD) cycle at 30 ℃ for 14 days shows that the frequency of locomotion increases during the dark phase. (B, C) Mean locomotion for 12 h (B) and 1 h (C) indicated that the locomotion significantly increases during the dark phase.

### Feeding activity

The diurnal rhythm of the feeding activity in crickets in a free-feeding environment was evaluated by counting the frames in which the head position overlapped with the feeding area (Fig 2). Due to the limited number of feeding events, it was not possible to generate double-plotted actograms and periodograms. There was no significant difference in feeding activity between the dark and light phases (Fig 4A). Nevertheless, acrophases of daily feeding activity rhythms clearly peaked around the transition from the light to the dark phase (Fig 4B, Fig S2A). Three out of six individuals showed an increase in feeding activity just before switching from dark to light, and their activities remained at a peak immediately after switching to light (Fig 4B). The other three animals exhibited a peak in feeding activity either immediately before or after the switch from light to dark (Fig 4B). Increased feeding activity during the transition between light and dark phases has also been reported in *Drosophila melanogaster* (Seay and Thummel, 2011; Ro et al., 2014; Niu et al., 2021). In *D. melanogaster*, feeding activities are concentrated near the lights-on, lights-off, or both (Seay and Thummel, 2011; Ro et al., 2014; Niu et al., 2021), possibly reflecting differences in the optimal timing of feeding activity between species. These results show that the quantitative analysis of feeding activity rhythms can be achieved by applying the prediction of body part position using DeepLabCut.

**Fig. 4.**
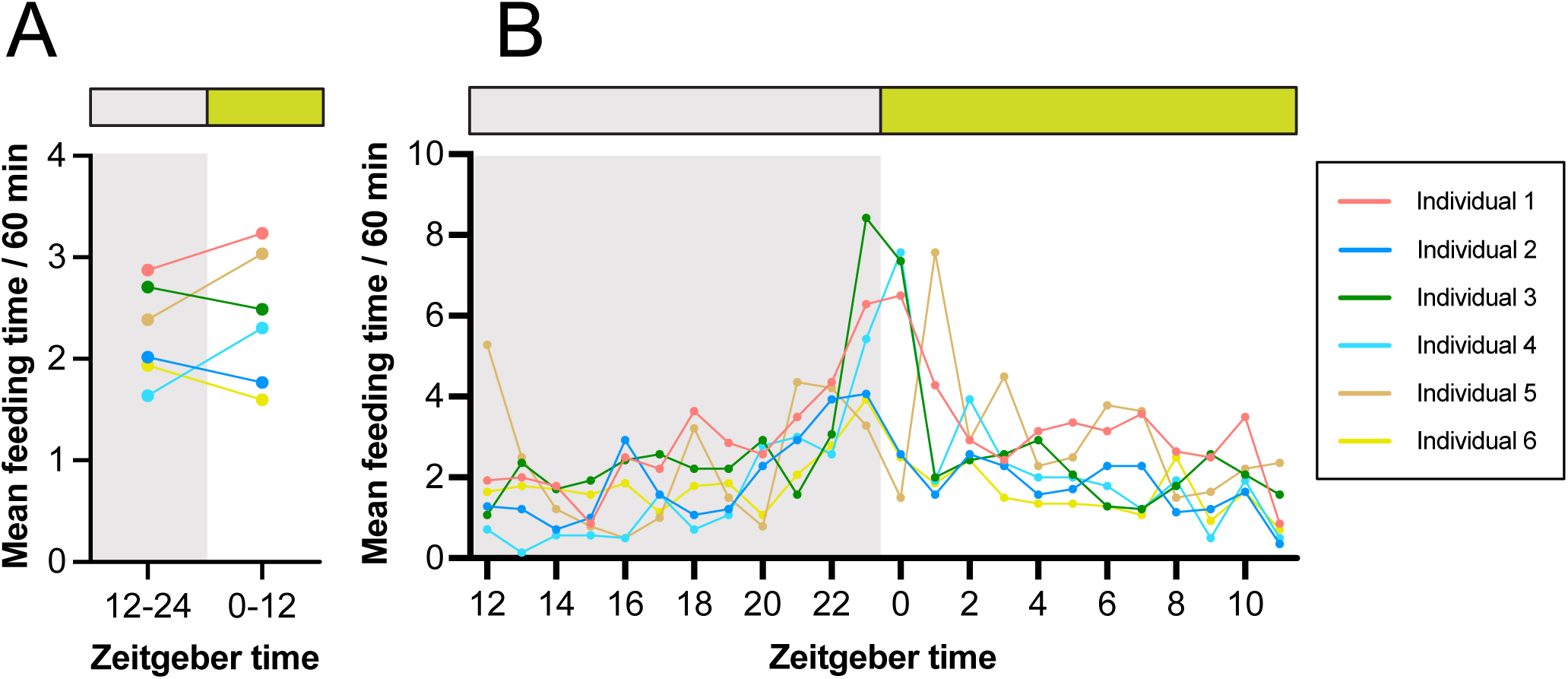
A quantitative analysis of feeding activity of crickets. (A, B) Mean feeding time for 12 h (A) and 1 h (B) suggested that peaks were evident around the transition from light to dark, though there was no significant difference in feeding activity between the dark and light phases.

### Resting state

Next, diurnal rhythms in the resting state of crickets were quantified. According to the definition of the SLS in *D. melanogaster*, an immobility time of more than 5 min was analysed in crickets (Fig 2). In contrast to the diurnal rhythm of locomotor activity, greater immobility was observed during the light period (Fig 5A, B). For most animals, the acrophases of the daily immobility time rhythm peaked several h after the transition from the dark to the light phase (Fig S2A). Consistent with the locomotor activity cycles, the rhythm of immobility time was also entrained to nearly 24 h (1440 min) by the 12:12 LD photocycle (Fig S2B). We also observed a bimodal rhythm in which a peak of immobility time was noted just before the end of the dark phase, in addition to the peak after lights-on (Fig 5C).

**Fig. 5.**
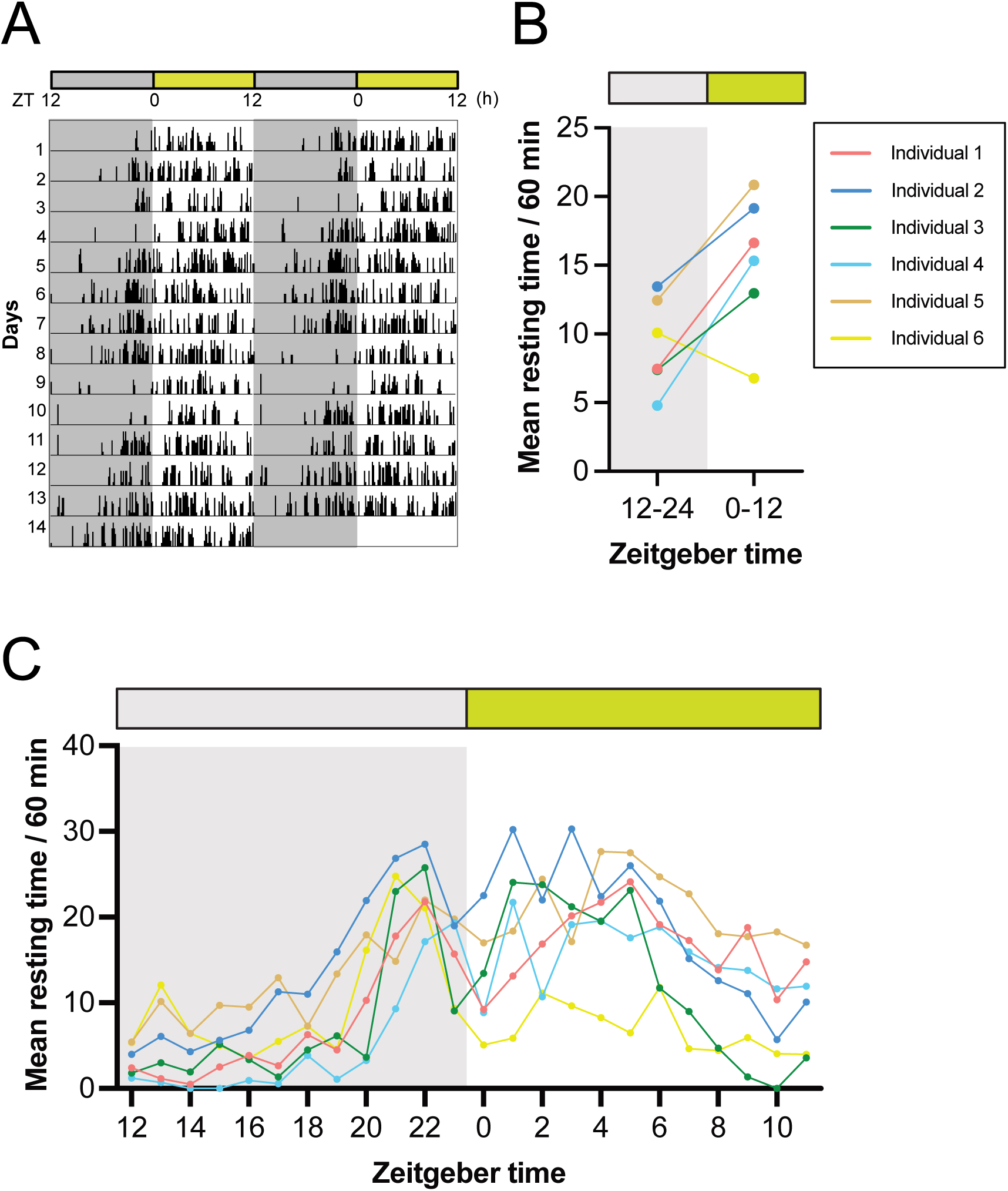
A quantitative analysis of resting time of crickets. (A) The double-plotted actogram recorded under a 12-h light:dark (LD) cycle at 30 ℃ for 14 days shows that the frequency of resting time increases during the light phase. Mean resting time for 12 h (C) and 1 h (D) indicated that the resting time significantly increases during the light phase, with a second peak in the late dark phase.

The behaviour-tracking system utilising DeepLabCut is particularly suitable for strictly defining the animal SLS based on a well-characterised sleeping posture (Campbell and Tobler, 1984). During the active phase, crickets support their bodies using the skeletal muscles, while during the resting phase, their hind legs are relaxed to lying position along with the antennae drooping (Fig S3A). This characteristic position may be associated with the SLS in crickets (Nishio, 2009). To quantify the diurnal rhythm of the resting state based on hind leg relaxation, we analysed the angle formed by three points from the coordinates of the abdomen, left hind legs, and right hind legs, which were predicted using DeepLabCut for each frame (Fig S3A). Four out of the six individuals showed a smaller hind leg angle in the light phase than in the dark phase, suggesting leg relaxation during the former phase (Fig 6A, B). To investigate the correlation between hind leg angle and immobility time, we performed the following two analyses. First, we calculated the average angle of the hind legs and the average immobility time for the same 1-h zeitgeber time over 14 days, investigating their correlation. The results revealed a negative correlation between the hind leg angle and immobility time in all individuals (Fig 6C, Fig S3B). Second, the average angle of the hind leg was calculated for immobility durations of 0, 1, 2,…, up to 20 min per individual. The results clearly show that the longer the continuous immobility time, the smaller the hind leg angle (Fig 6D, Fig S3C). These data suggest that hind leg angle is strongly negatively correlated with an individual’s immobility time and that postural assessment based on hind leg angle may be useful for estimating SLS more precisely.

**Fig. 6.**
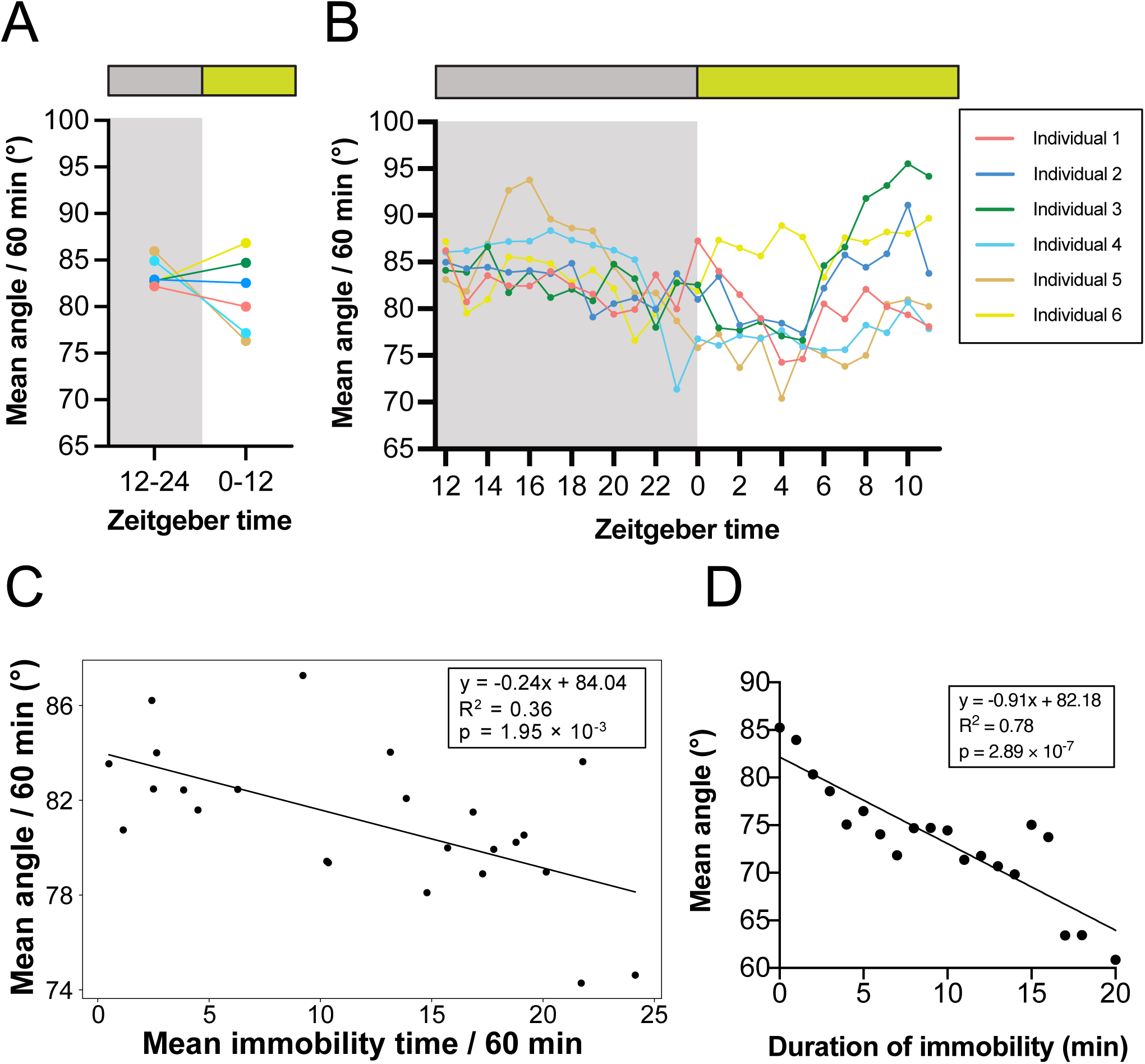
Body posture associated with the resting phase in crickets. The body posture and leg relaxation were calculated from the angle between the three points of abdomen, left hind leg, and right hind leg. The relationship with resting was then examined. (A, B) The mean angle for 12 h (A) and 1 h (B) indicated that leg relaxation was significantly increased during the light period, when the frequency of resting was higher. (C) A scatter plot of the mean angle per h against the mean resting frequency corresponding to same h was calculated with a fitted linear regression line. The equation of the line, with R squared as the coefficient of determination and p-value, indicating the negative correlation between hind legs angle and immobility time. (D) Mean hind legs angle was calculated for each immobile time from 0 to 20 min with a fitted linear regression line. The equation of the line, with R squared as the coefficient of determination and p-value, indicating a strong negative correlation.

## Discussion

In this study, we developed the system using the DeepLabCut to automatically analyse the diurnal rhythms of behaviors and postures of crickets, which are a well-established model organism for studies into the circadian rhythm and behaviour of hemimetabolous insects. Our system enabled us to simultaneously measure the rhythms of locomotive activity, in addition to feeding behaviour and SLS rhythms in a single experimental run. DeepLabCut has provided analysis of the diurnal rhythm of SLS by postural estimation based on hind legs, as well as by the conventionally adopted criterion of at least 5 min of continuous immobility time in *D. melanogaster* (Shaw et al., 2000; Hendricks et al., 2000). Our system is expected to save researchers a great deal of time, generate reproducible results by eliminating inter-researcher variability, and enable experiments that are not possible with other methods, such as the seesaw-type actometer (Moriyama et al., 2008) and the infrared sensor-based actometer (Yamano et al., 2001).

Our study successfully leveraged DeepLabCut to automatically label cricket body parts with high accuracy, but low-confidence labels (likelihood < 0.95) were included at approximately 5% or less during the light phase and at about 5-15% during the dark phase. Although the arena for behavioural experiments used in this study was designed to be user-friendly and easy to set up, low-confidence labels were believed to be generated by the reflection of light from LEDs on the acrylic case and by the shadows cast by objects in the acrylic case. In this study, low-confidence labels with likelihood < 0.95 were treated as missing values and supplemented with the mean values of the pre- and post-frames. The environment setting and number of training sessions of the model should be determined to ensure the spatial resolution required to detect the behaviour and posture of interest to researchers.

In this study, we applied the SLS criterion used in *D. melanogaster*, “continuous immobility for at least 5 min,” to crickets and were able to quantify the diurnal SLS rhythms. Moreover, we were able to quantify hind leg relaxation based on the positional coordinates predicted by DeepLabCut, allowing us to analyse the SLS in a different manner than in previous studies. Immobility time was observed at high levels during the light phase but also showed an additional transient peak in the late dark phase, consistent with previous studies in Drosophila (Dubowy and Shegal, 2017). Analysis of the SLS rhythm focusing on hind leg relaxation showed that the average angle of the hind legs was smaller in many individuals during the light period. Notably, there was a decrease in the average angle of the hind legs in the late dark phase but only in approximately half of the individuals used in this experiment, suggesting that crickets may have had short bouts of siesta sleep during the dark phase (active phase). Thus, it is possible that resting behaviours observed during the late dark phase and those observed during the light phase (inactive phase) may have different physiological significance. Future investigation would require careful observation of individuals at a fine temporal resolution, focusing on movements such as posture, tactile sensation, and digestion.

In addition, there was a strong negative correlation between hind leg angle and immobility duration, suggesting that the hind leg angle can be a reliable indicator for detecting SLS. However, the thresholds for homeostasis, arousal, and immobility duration, which are characteristics of the SLS, may differ among species (Allada et al., 2017; van et al., 2013). Furthermore, it is helpful to perform electrical brain measurements to verify whether the observed SLS is true sleep. In the future, homeostatic assessment and electrophysiological approaches will be required to advance sleep research in crickets.

In summary, we report the DeepLabCut-based accurate detection of multiple behaviours and postures over two weeks in multiple crickets. Our system can be easily applied to various animal species other than crickets, and allows for the evolutionary study of many physiological processes, including sleep. By combining high-fidelity behavioural tracking with genomic and transcriptomic data, it is possible to explore and understand the molecular basis of behavioural variations and phenotypic plasticity.

## Materials and Methods

### Animals

All experiments were performed with adult virgin female crickets (*Teleogryllus occipitalis*) that were collected in Amami Oshima Island, Kagoshima, Japan and cultured in a laboratory, maintained under standard environmental conditions of 12 h light and 12 h darkness (LD 12:12, light: 0600–1800 h; Japan Standard Time) at a constant temperature of 30 ℃. The animals were provided with cricket food (Tsukiyono Farm, Gunma, Japan) and water *ad libitum*. In the recordings, six individuals were placed individually in their cases, and all six were simultaneously recorded. The reason for using virgin females in this study is to prevent chirping sounds produced by adult males from influencing others.

### Recording system

The recording system used in this study is shown in Fig 1. The crickets were individually placed in transparent acrylic cases (150 × 100 × 50 mm). To render the individuals invisible to each other, sheets of paper with a height of 15 mm were placed between each acrylic case. Imaging was conducted using an infrared camera module (Raspberry Pi NoIR; 3,280 × 2464 pixels) connected to a Raspberry Pi 3 model B (Raspberry Pi Foundation, Cambridge, UK), which captured RGB images of the crickets at 1-min intervals. The camera was positioned 450 mm above the bottom of the acrylic case. The experimental system was installed inside an incubator with an air temperature maintained at 30 □, and humidity was kept at 45 ± 5%. Illumination was performed using white LEDs (Timely, Tokyo, Japan) and infrared LEDs (Broadwatch, Tokyo, Japan). The white LEDs were connected to a time switch, ensuring a 12-h light and a 12-h dark period (LD 12:12; Light period: 6:00 to 18:00, Japan Standard Time; 0 to 12, Zeitgeber Time). Light intensity at the position of the animals was measured using a spectroradiometer (MS-720; Eko Instruments, Shibuya, Tokyo, Japan) and was approximately 150 lux. After entraining the crickets to a cycle of LD 12:12 light cycle for 3 days (not recorded), 14-day time-lapse imaging was conducted. During this period, food and water replenishment and replacement were carried out three times, at intervals of 3 and 4 days, respectively.

### Data acquisition

All images obtained were cropped such that each acrylic case for each individual fit (Fig 2). A series of images were created as a single movie and fed into DeepLabCut. The lens distortion, which depends on the distance and angle between the camera and the subject, was evaluated based on 10 mm × 10 mm grid squares placed under the acrylic cases. The maximum difference in length owing to lens distortion at the centre point and the four corners of the image was less than one pixel. This corresponded to 0.68% of the body length of the crickets used in this experiment. Furthermore, this was smaller than the average offset from automatic labelling via machine learning, as described later. Therefore, it was concluded that lens distortion in this recording environment did not affect behaviour quantification, and correction for lens distortion was not performed.

### DeepLabCut training and evaluation

We used DeepLabCut for the MacOS M1 chip (*i.e.*, DEEPLABCUT_M1) to quantify cricket locomotion, feeding activity, immobility time, and posture. The aim was to construct a model that could automatically estimate six points on the body of a cricket: the head, thorax, abdomen, abdominal tip, left hind leg, and right hind leg (Fig 2). To generate training datasets for the first training, 120 frames were randomly sampled from each video using k-means clustering in DeepLabCut. Six points on the body of the cricket were manually labelled. In the first training step, a ResNet-50-based neural network with 50,000 iterations was used. Because the model performance was initially suboptimal, we refined the model after the first training. During refinement, the keypoints labelled in 120 randomly extracted frames were manually corrected. These training and refinement processes were repeated four times, culminating in five training sessions to construct the final model. The number of iterations was set to 50,000 from the first to third sessions, 200,000 from the fourth session, and 500,000 from the fifth session.

To evaluate the training of the model, the relationship between the iteration number and the loss value was calculated. Next, the mean deviation (px) of the coordinates of the six keypoints between the manually labelled teacher data and automatic labelling by the model was calculated for each training set as a training error. Simultaneously, the mean deviation (px) of the coordinates of the six keypoints between the manually labelled data without the teacher data and automatic labelling was calculated for each training set as a test error. Additionally, the proportion of data with a likelihood value of 0.95 or higher and less than 0.95 as missing values, representing the estimated probability for each keypoint, was calculated to assess the labelling accuracy across all data. When the keypoints of a cricket were obscured by another object, the model attempted to approximate its position. The likelihood value of such frames is typically low. In this study, coordinates with a likelihood value of less than 0.95 were removed. The missing values were imputed by mean imputation using a Python script (Fig 2).

### Behaviour analysis

Locomotor activity, feeding activity, and SLS were defined for each frame based on the coordinates of the keypoints of body parts (*i.e.* head, thorax, abdomen, abdominal tip, left hind leg, and right hind leg) (Fig 2). Locomotor activity was defined as the movement of a distance equal to or greater than body length within one frame (1 min). Feeding activity was defined when the head coordinates were located within the feeding area. SLS was defined as a change in the distance of the abdomen’s coordinates within one frame (1 min) of less than three pixels and when this immobile state persisted for five or more frames (equivalent to five or more minutes). Criteria for > 5 min of immobility were adopted from the definition of SLS in fruit flies (Shaw et al., 2000; Hendricks et al., 2000). Food and water were replenished and replaced thrice during the 14 days of recording. As the position of the acrylic case shifts before and after these replenishments, the coordinates of the feed area were adjusted accordingly.

Raw data were visualised as double-plotted actograms using ActogramJ (Schmid et al., 2011). Periodicity analysis was conducted using the Chi-square periodogram (Sokolove and Bushell, 1978) and the LombScargle periodogram (Ruf, 1999). These approaches were selected to consider their characteristics and the risks of methods (Tackenberg & Hughev, 2021). If a peak in the periodogram exceeded a confidence level of 0.05 (α = 0.05), it was considered as a significant rhythm (Kaneko et al., 2000). Oriana software, v.4 (Kovach Computing Wales, UK) was used to calculate the average acrophase (peak frequency of a certain behaviour within 24 h) during the experimental period (Kovach, 2011). To compare the levels of behaviours or posture (*i.e.* locomotor activity, immobility time, and hind leg angle) during the light and dark periods, or for the analysis of hourly trends at each level, hourly averaged data were used. For feeding activity, the averages for every h and every 12 h were used because the total activity was small.

### Quantification of the angle between the abdomen and both hind legs

The coordinates of the abdomen, left hind legs, and right hind legs were used to calculate the angles hind leg opening. A Q-Q plot was used to check whether the angle datasets followed a normal distribution. The top and bottom 2.5% of the angle data were considered outliers and excluded, with the data within the 95% confidence interval used for further analysis. Postures in which the angles of leg opening were outliers included those in which the crickets brought their hind legs up to their mouths. To statistically assess the difference in the angle during the dark and light phases, the angles were averaged every h, and the change over a day was calculated every h and every 12 h. To investigate the relationship between the angles and immobility time, the linear regression correlation coefficient was computed between the angle and the immobility time for each h.

### Statistics

Student’s *t*-test were used to compare differences in the means of activities between night and day. Two-way analysis of variance (ANOVA) followed by Sidak’s multiple comparison test was used to test for differences in behavioural transition probabilities. The Shapiro-Wilk test and the Kolmogorov-Smirnov tests were used to determine whether the angle data followed a normal distribution. Linear regression analysis was used to demonstrate the correlation between the SLS and the angle between three points: the abdomen and hind legs. Periodic and circular analyses were performed as described previously. Statistical significance was set at *P* < 0.05.

## Acknowledgments

This work was supported in part by the Cabinet Office, Government of Japan, Cross-ministerial Moonshot Agriculture, Forestry and Fisheries Research and Development Program, and “Technologies for Smart Bio-industry and Agriculture” (funding agency: Bio-oriented Technology Research Advancement Institution) (JPJ009237). We would like to thank Editage (www.editage.jp) for English language editing.

## Author contributions

S.H. designed the study, the main conceptual ideas, and the proof outline. S.H. collected the data. K.K., M.Y., T.A and T.S. aided in interpreting the results and worked on the manuscript. T.A. and T.S. supervised the project. S.H. wrote the manuscript with support from K.K. and T.S. All authors discussed the results and commented on the manuscript.

## Figure legends

**Fig. S1. Model accuracy evaluation.** (A) The loss value was calculated based on the discrepancies between the model’s predicted keypoint coordinates and the actual labelling data. This graph shows that the loss value decreases as the number of iterations increases. (B) The train error is calculated as the difference in distance (px) between the actual labelled coordinates and the model’s predicted keypoint coordinates, averaged over 6 keypoints and the number of teacher data (black circles). Test error was calculated in the same way as Train error, using a dataset reserved for validating the model’s performance (red circles). These graphs show the accuracy improved with an increased number of training iterations. (C) Example images of the train error by the model after the 5th training (manual: circles, automatic: crosses). (D) The percentage of labelling accuracies was calculated as the percentage of frames for which the likelihood of validity that was greater than or equal to 0.95 for each coordinate of the output raw data (N=6). (E) The missing value was calculated as the percentage of frames for which the likelihood was less than 0.95 and was indicated by light and dark phases for each keypoint (N=6). The error bars indicate standard error of the mean (SEM).

**Fig. S2. Behavioural profile of each of the six individuals** (Colours correspond to individual numbers). (A) Acrophase for locomotion, feeding behaviour, and resting state were calculated by angle mean and vector length weighting from circular analysis, respectively. The red line shows the median of each behaviour. (B) The periodograms for locomotion and resting state were created by Chi-square and Lomb-Scargle statistics. This graph demonstrated that the frequencies of both locomotion and resting state have a circadian rhythm of approximately 24 h (1440 min), whereas individual #6 had a cycle shorter than 24 h.

**Fig. S3. The body posture associated with resting phase in crickets.** (A) Postural features at awake (the left panels) and resting (the right panels) states were captured from the side (the upper panels) and from above (the lower panels). The decrease in the height of femur and that of the abdomen, leg relaxation (calculated based on the angle between the three points of abdomen, left hind leg, and right hind leg) suggested that these postures are characteristic of the resting. (D) A scatter plot of the mean angle per h against the mean resting frequency corresponding to same hour was calculated with a fitted linear regression line in each individual. The equation of the line and Pearson’s r value are provided, indicating a relatively strong negative correlation, with variation among individuals. (E) The mean angle was calculated for each immobility time from 0 to 20 min with a fitted linear regression line. The equation of the line, with R squared as the coefficient of determination, is provided, indicating a relatively strong negative correlation in each individual.

## Movie

**Movie. Labelled video output from DeepLabCut.** Automatic labelling by the model after the fifth training showed high accuracy for the six keypoints of the crickets in both the dark and light phases, independent of the LEDs and objects.

