## Supplementary figures and images for "DeepLabCut-based Behavioural and Posture Analysis in a Cricket"

### Supplemental Figure 1

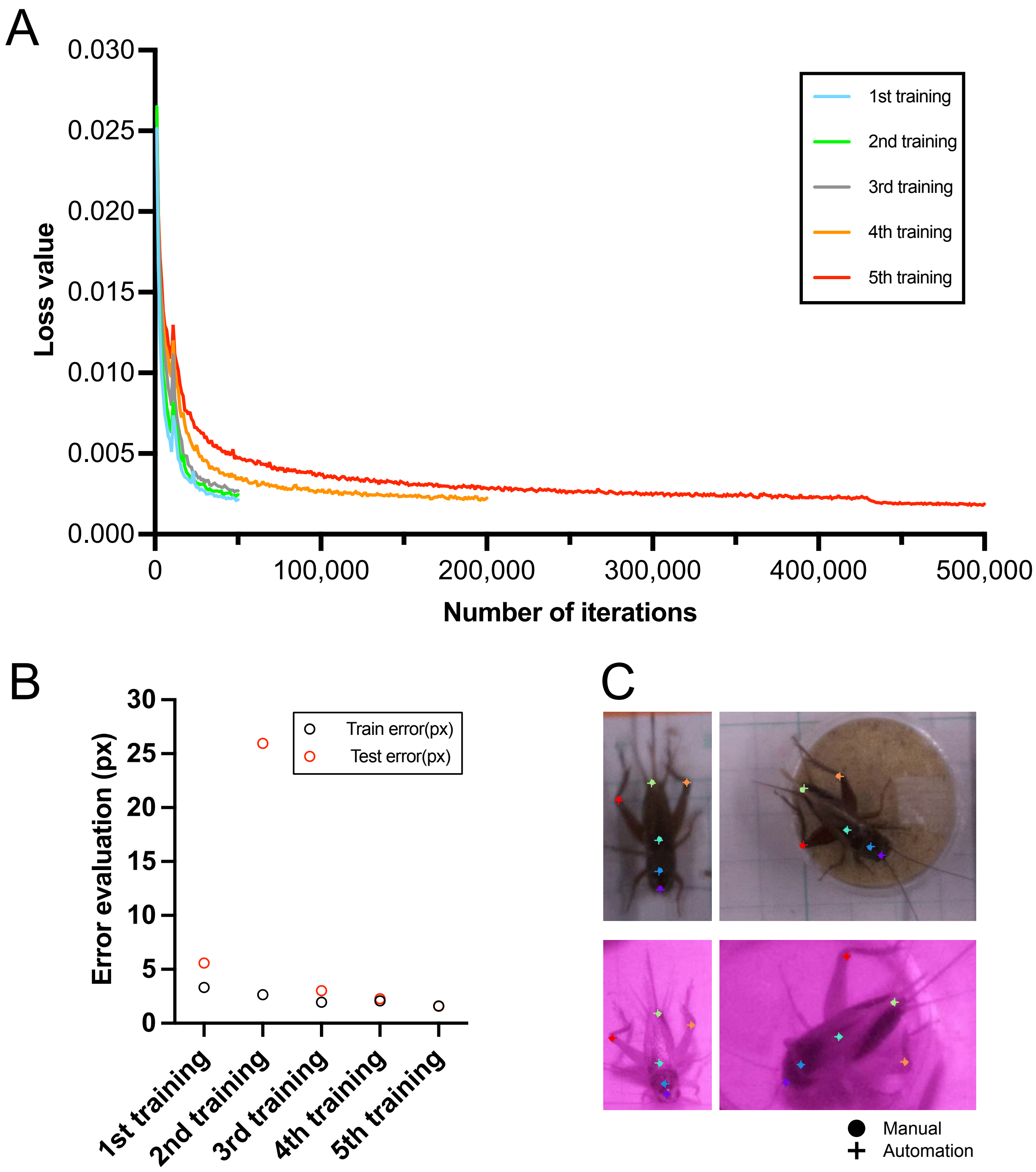

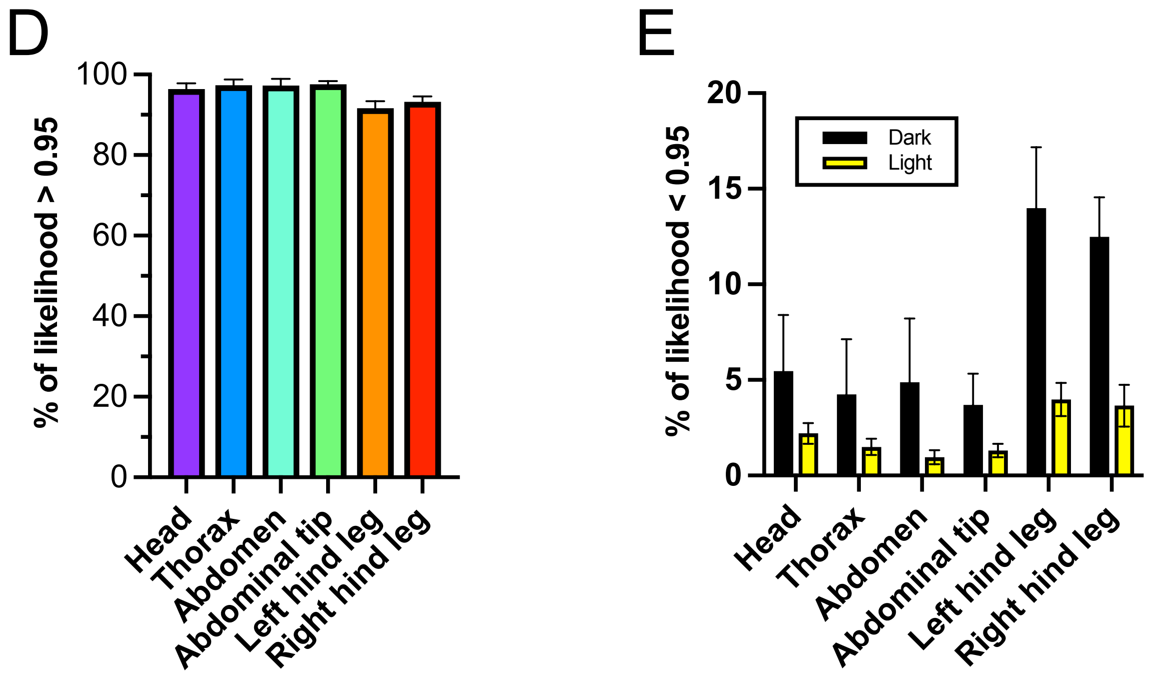

### Supplemental Figure 2

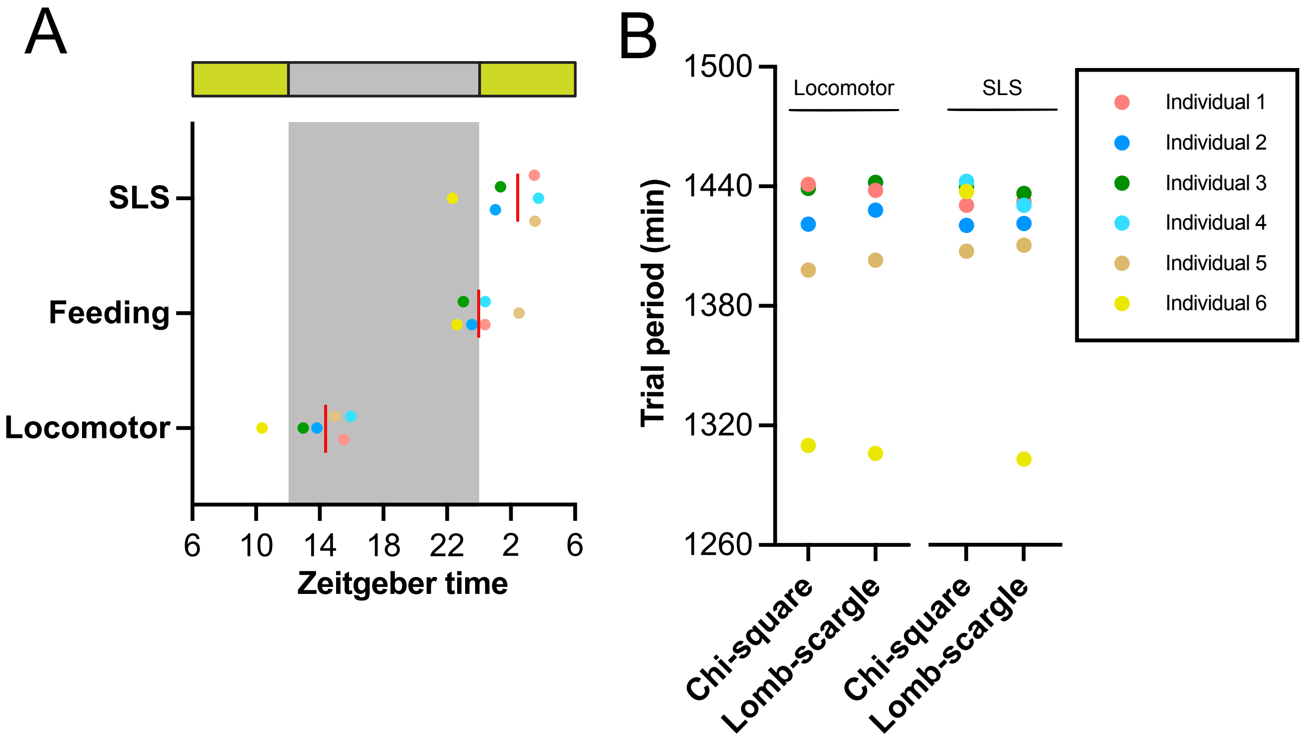

### Supplemental Figure 3

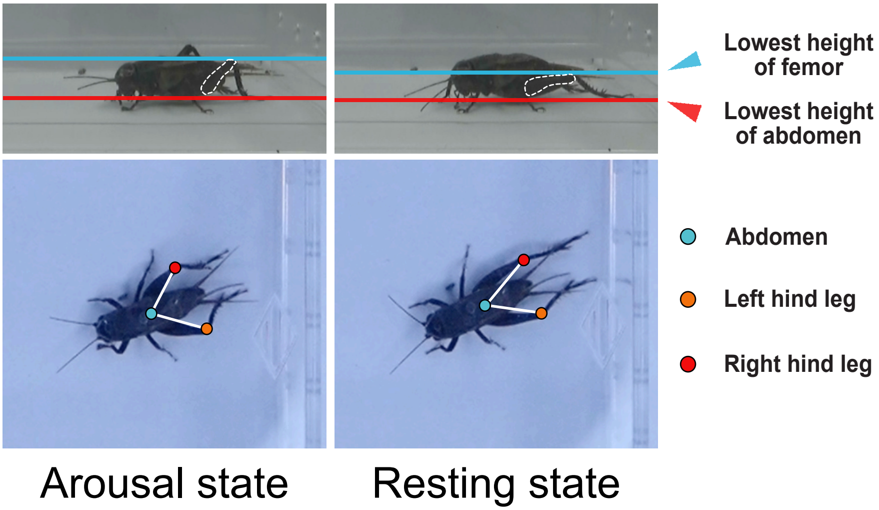

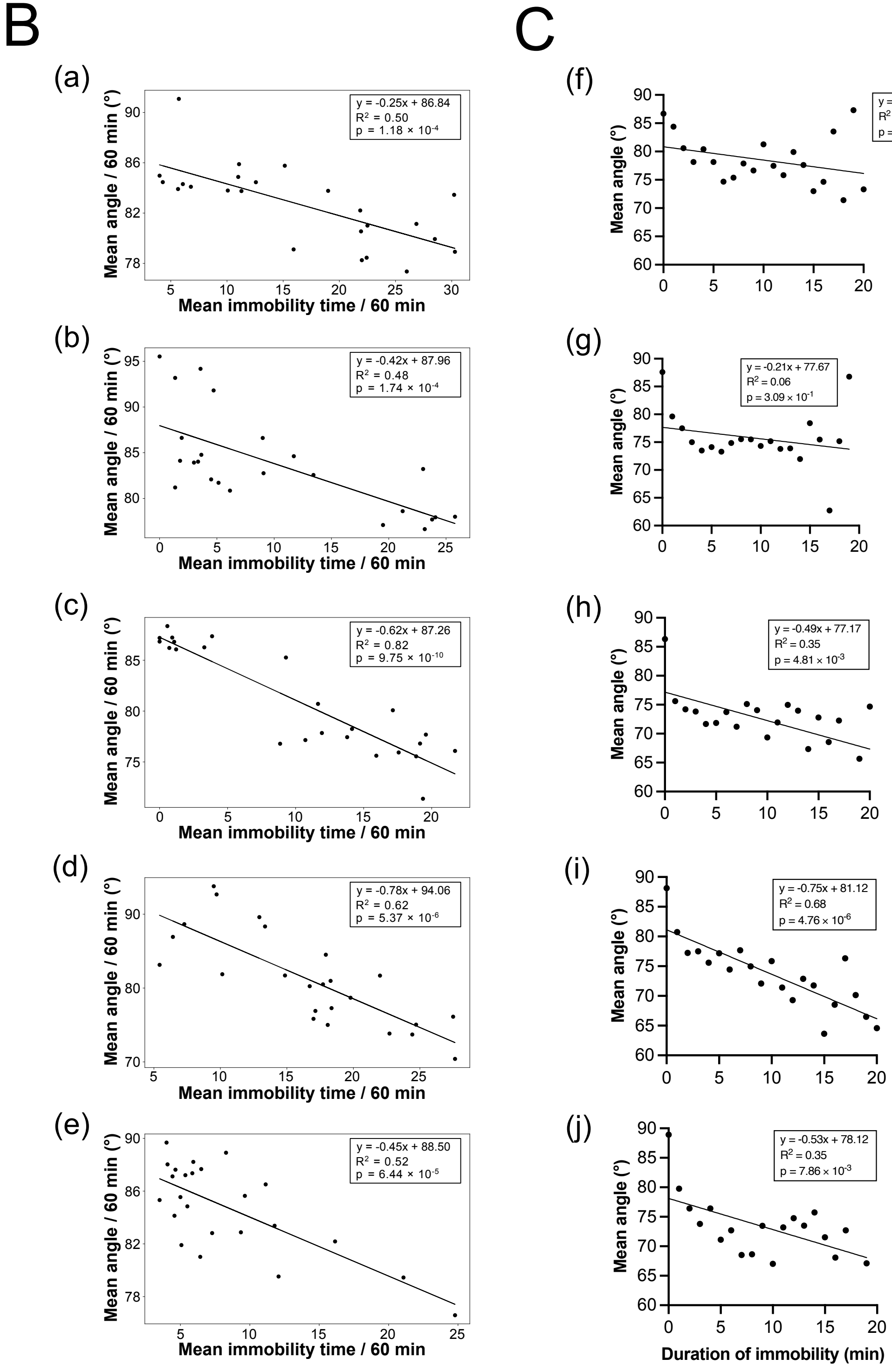
